# Cell type mapping of mild malformations of cortical development with oligodendroglial hyperplasia in epilepsy using single-nucleus multiomics

**DOI:** 10.1101/2024.12.12.628140

**Authors:** Isabella C. Galvão, Manuela Lemoine, Ludmyla Kandratavicius, Clarissa L. Yasuda, Marina K. M. Alvim, Enrico Ghizoni, Ingmar Blümcke, Fernando Cendes, Fabio Rogerio, Iscia Lopes-Cendes, Diogo F. T. Veiga

## Abstract

**Objective:** Mild malformations of cortical development with oligodendroglial hyperplasia in epilepsy (MOGHE) are brain lesions associated with focal epilepsy and characterized by increased oligodendroglial density, heterotopic neurons, and hypomyelination in the white matter. While previous studies have implicated somatic mutations in the *SLC35A2* gene, the cellular and molecular mechanisms underlying MOGHE pathogenesis remain elusive. To address this gap, this study aimed to systematically characterize the cell type composition and molecular alterations of MOGHE lesions at cellular resolution using single-nucleus multiomics profiling.

**Methods:** We performed single-nucleus multiomics sequencing to obtain paired gene expression and chromatin accessibility profiles of > 31,000 nuclei from gray matter and white matter regions of MOGHE lesions, and compared the results with publicly available neurotypical control datasets.

**Results:** The analysis of gray and white matter regions from two MOGHE patients revealed significant cellular composition alterations, including the presence of heterotopic neurons and disease-specific oligodendrocytes populations within the subcortical white matter. MOGHE-specific oligodendrocytes were characterized by the upregulation of synaptic functions and enhanced neuron communication, denoting a possible role in synaptic support and the mediation of glial-neuron interactions in the disease. On the other hand, MOGHE heterotopic neurons were characterized by the upregulation of genes associated with neuronal migration and the Wnt signaling pathway, suggesting a mechanism underlying their atypical localization.

**Significance:** This high-resolution cell type mapping of MOGHE lesions in clinical samples unveils neuronal and glial populations affected by the disease, and provides novel insights into the pathophysiological mechanisms of MOGHE.

**Key Points:**

- We provide a multimodal cellular atlas of the human cortical and subcortical regions affected in MOGHE
- MOGHE-associated oligodendrocytes showed upregulation of synaptic functions and enhanced neuron communication
- Neuronal migration and Wnt signaling are upregulated in MOGHE heterotopic neurons

## INTRODUCTION

Malformations of cortical development (MCDs) encompass a varied group of disorders that arise due to disruptions in the development of the cerebral cortex. MCDs can affect individuals across all age groups, although symptoms typically begin in early childhood and extend to young adulthood and may include epilepsy, developmental delays, and intellectual impairments^1^. MCDs typically manifest as focal epilepsy and include focal cortical dysplasia (FCD) as well as mild malformations of cortical development with oligodendroglial hyperplasia in epilepsy (MOGHE), a neuropathological entity featuring abnormalities predominantly in the white matter, while maintaining normal cortical lamination^2^. This latter characteristic distinguishes MOGHE from FCD, which is characterized by abnormal cortical architecture associated or not with abnormal cell types^3^. MOGHE was proposed as a distinct pathological entity in 2017^2^, and was recently included in the International League Against Epilepsy (ILAE) classification of cortical malformations^4^. The increase in oligodendroglia and the appearance of heterotopic neurons in the white matter, together with the blurring of the gray-white matter junction, are the neuropathological hallmarks of this condition. The affected white matter tissue presents with proliferation of oligodendroglial cells expressing Olig2 and irregular areas of hypomyelination due to reduced myelin protein expression. Clinically, MOGHE manifests as focal epilepsy and treatment might require surgical resection of the affected region. MOGHE lesions occur most often in the frontal lobe and have early seizure onset, typically in young children, but may also occur in the temporal lobe, and symptoms may manifest later even in adulthood^5,6^.

Genetic investigations have found that MOGHE patients frequently harbor somatic brain variants in the *SLC35A2* gene, which encodes a UDP-galactose transporter^7^. One study reported that 45% (9/20) of MOGHE cases presented truncating or frameshift variants in *SLC35A2*^7^, while another screening identified *SLC35A2* variants in 47.1% (16/35) of cases^8^. In agreement, the loss of *Slc35a2* has been shown to recapitulate the MOGHE-like phenotype in conditional knockout mouse models^9–11^. Other genes may also play a role, as demonstrated by a case identified with a variant in the *NPRL2* gene^8^. However, the cellular and molecular mechanisms contributing to MOGHE pathogenesis have not been systematically studied.

Single-nucleus RNA sequencing (snRNA-seq) and single-nucleus assay for transposase accessible chromatin sequencing (snATAC-seq) have been applied to dissect the cellular diversity of brain tissues in neurological conditions, enabling the discovery of dysregulated cell states involved in diseases including focal MCDs^12–16^. For instance, we previously applied paired snRNA-seq and snATAC-seq to identify cell populations involved in FCD type II pathology, which allowed us to uncover vulnerable neuronal subtypes and activated microglial states in these cortical malformations^15^.

In this study, we employed multiomics single-nucleus profiling to systematically analyze the cellular landscapes of both gray matter and subcortical white matter of MOGHE tissues, and compared the results with publicly available control datasets. As a result, we characterized perturbed oligodendroglial and neuronal cell populations associated with the condition. Our findings provide novel insights into the pathophysiological mechanisms of MOGHE.

## MATERIALS & METHODS

### Clinical samples and neuropathological diagnosis

Fresh brain samples were collected from two patients who underwent surgery for drug-resistant focal epilepsy at the Hospital de Clínicas, University of Campinas. All procedures were approved by the Research Ethics Board of our Institution (CAAE: 12112913.3.0000.5404), and written informed consent was obtained from patients or their legal guardians before surgery.

Representative surgical specimens were either formalin-fixed and paraffin-embedded (FFPE) or snap-frozen in liquid nitrogen and stored at −80°C. FFPE samples were submitted to routine diagnostics in serial 4 µm-sections stained with hematoxylin and eosin (H&E) and submitted to immunohistochemical reactions. For the latter protocol, the sections were exposed to antibodies against Olig2 (glial marker; dilution 1:250, polyclonal, Millipore Sigma, cat#AB9610, Darmstadt, Germany), and MAP2 (neuronal marker; 1:1,000, clone M13, Thermo Fisher, cat#13-1500, Waltham, MA, USA), overnight at room temperature. Then, a detection solution containing the secondary antibody and peroxidase (AdvanceTMHRP®, Dako, cat#K4068, Glostrup, Denmark; or EnvisionTM Flex+, Dako, cat#K8002, Glostrup, Denmark) was added for 30 min at 37°C. 3,3-diaminobenzidine (DAB) was used as a chromogenic substrate and counterstaining was performed with hematoxylin. Negative controls (without primary antibody) were run concurrently with all immunohistochemical reactions. Digitized images were obtained using a CS2 Aperio ScanScope scanner (Aperio Technologies, Vista, CA, USA).

Samples were neuropathologically classified as MOGHE according to the latest guidelines of the ILAE^4^. Specifically, the specimens showed an increase in oligodendroglial cells (> 2200 Olig2-positive cells/mm^2^) and an excess of heterotopic neurons in the white matter (>30 MAP2-positive neurons/mm^2^). No cortical cytoarchitectural changes were observed. After histopathological confirmation of MOGHE, a mirror-frozen section of the tissue used for FFPE analysis was dissected using a surgical scalp in a Petri dish on dry ice, allowing separation of the GM and WM. Subsequently, tissue samples were submitted to molecular analyses as described below.

### Whole-exome sequencing and somatic mutations

Genomic DNA was extracted from MOGHE-WM tissue using the PureLink Genomic DNA Mini Kit (cat# K182001, Invitrogen) following the recommended protocol. Exome capture libraries were generated using the SureSelect Human All Exon v8 kit (Agilent). Paired-end sequencing was performed at 101 base pairs on each side of the DNA fragment on the Illumina NovaSeq 6000 platform. Deep sequencing obtained an average of 489 M reads per sample, with a mean depth exome coverage of 530x.

Reads were aligned to the hg38 human reference genome using bwa mem 0.7.17^17^. Bam files were sorted and duplicates were removed using Picard tools v.2.18.2. Indel realignment and base quality score recalibration were performed with GATK^18^. Variants were called using the GATK haplotype caller v4.0.5.1^18^ and annotated using SnpEff v.4.3^19^. Somatic mutations including single-nucleotide variants and indels were detected using Mutect2 (GATK v.4.0.5.1) with the 1000 Genomes panel of normals as controls.

### Nuclei isolation and multiome library construction

Nuclei were isolated using the Chromium Nuclei Isolation kit (10X Genomics) according to the manufacturer’s protocol (CG000505, revision A), which is compatible with the Single Cell Multiome ATAC + Gene Expression assay. Briefly, frozen tissue samples were dissociated with a pestle in lysis buffer and passed through a nuclei isolation column. The isolated nuclei were then resuspended in a debris removal buffer, followed by washes to clear any remaining debris. Recovered nuclei were stained with Trypan blue and counted with a hemocytometer. Nuclei integrity was evaluated under a microscope at 40X or 60X magnification, and nuclei were considered viable if they were roundish and with an intact membrane. We then performed library construction, which included open chromatin transposition and droplet (GEM) formation using a Chromium Controller (10X Genomics). Libraries were prepared using the Multiome ATAC + Gene Expression kit (10X Genomics) following the manufacturer’s protocol (CG000338, Revision E). The quality of the purified ATAC and RNA libraries was assessed using TapeStation (Agilent Technologies).

### Sequencing and raw data processing

Multiome libraries were sequenced on the Illumina NovaSeq 6000 platform according to the sequencing depth and read length guidelines specified in the Multiome kit protocol (CG000338). Demultiplexing, genome alignment, gene quantification, and peak accessibility analyses for single nuclei were conducted using the Cell Ranger ARC pipeline. Specifically, fastq files were generated with cellranger mkfastq, followed by processing with cellranger-arc count v.2.0.2 using the human genome reference GRCh38 to produce count matrices for RNA and ATAC modalities. Downstream data processing for both snRNA-seq and snATAC-seq were performed using Seurat v.5.0.1^20^ and Signac v.1.12.0^21^ in R v.4.1.2.

### snRNA-seq analysis

Ambient RNA decontamination was carried out using DecontX^22^ as implemented in the celda^23^ v.1.10.0 R package. Briefly, RNA counts for each sample were processed with the decontX function, utilizing the raw droplet matrix generated by cellranger count as background. DecontX then employs a Bayesian approach to estimate and remove contamination in individual cells. Nuclei were filtered using Seurat based on the following criteria: 1000 < UMI counts < 25,000, nFeatures > 400, and mitochondrial read percentage (percent.mt) < 15. Doublet identification was conducted for each sample using scDblFinder^24^ v.1.8.0.43.

Standard Seurat processing and normalization steps were applied in each sample, including NormalizeData, FindVariableFeatures, ScaleData, RunPCA, RunUMAP, FindNeighbors, and FindClusters. The filtered counts matrix was log-normalized with regression to correct for mitochondrial gene percentage. To correct batch effects across samples, we recomputed NormalizeData, FindVariableFeatures and RunPCA on the merged object, and Harmony^25^ v.1.2 was used to integrate PCA projections with automatic hyperparameter optimization and default settings. Harmony’s algorithm projects cells into a shared space where they cluster based on cell type rather than sample-specific attributes like sequencing batch. Dimensionality reduction was performed using RunUMAP with the first 40 principal components identified using the elbow plot. Clustering was executed with FindNeighbors and FindClusters functions employing the smart local moving (SLM) algorithm, with a resolution set to 1.2.

Cell type markers were obtained by applying the Wilcox rank-sum test to RNA-normalized data, with a log2 fold-change threshold of 0.25 and an adjusted P-value of less than 0.05. Differentially expressed genes (DEGs) were identified using MAST. For identifying DEGs in MOGHE-WM vs Control-WM, MAST was applied using the number of counts, mitochondrial percentage and dataset as latent variables. For MOGHE-WM vs MOGHE-GM, we used the number of counts, mitochondrial percentage, donor, and brain region as latent variables. Genes with an adjusted P value of less than 0.05 and fold-change threshold of 2 were considered significant. Mitochondrial genes and those located in sex chromosomes were removed from the analysis.

Gene Ontology (GO) and Disease Ontology (DO) enrichment was assessed with clusterProfiler^26^ v.4.2.2 and DOSE^27^ v.3.20.1 R Packages using the enrichGO and enrichDO functions, respectively. Enrichment analysis with the Synaptic GO (SynGO) ontology^28^ to find overrepresented synaptic terms was performed in the SynGO online portal v. 1.2 (http://syngoportal.org) using the set of brain expressed genes as the background.

### Integration with snRNA-seq from heterotypical white matter

We incorporated control white matter snRNA-seq data from two autopsy donors with no neurological history, originally sequenced in Elkjaer et al.^29^. Raw sequencing data were obtained from GEO (accession number GSE231585) and processed with cellranger count v.7.0.1 using the human genome reference GChR38. Nuclei were filtered using Seurat based on the following criteria: 1000 < UMI counts < 25,000, 200 < nFeatures < 4000, and mitochondrial read percentage (percent.mt) < 1. This dataset was integrated into our MOGHE dataset using the procedure outlined in the “snRNA-seq analysis” section.

### snATAC-seq analysis

Samples were preprocessed in Signac to filter out low-quality nuclei based on these criteria: 1,000 < ATAC fragments < 50,000, ATAC fragments in peaks > 400, nucleosome signal < 2, and TSS enrichment > 2. Peaks within each sample were identified using MACS2^30^. Standard processing and normalization were applied with the following functions: RunTFIDF, FindTopFeatures, RunSVD, and FindClusters. The peak counts matrix for each sample was normalized using RunTFIDF to adjust for differences in sequencing depth. Dimensionality reduction was performed with latent semantic indexing (LSI) via the RunSVD function, with UMAP projection utilizing LSI components 2-30, as the first LSI component was determined to represent technical variation. Prior to sample integration, a common peak set across samples was created using MACS2, and peaks in each sample were quantified with the FeatureMatrix function. Peaks located in non-standard chromosomes and blacklisted regions were excluded from subsequent analyses. We then re-computed RunTFIDF, FindTopFeatures, and RunSVD on the merged dataset, and applied Harmony^25^ to integrate low-dimensional cell embeddings (LSI components 2-30) across samples, using parameters similar to those defined for RNA analysis. Chromatin accessibility at the gene level was computed with the GeneActivity function from the Signac package, which sums the fragments intersecting the gene body and promoter regions of each gene.

### Multimodal data integration

To integrate gene expression and chromatin accessibility data, we utilized Weighted Nearest Neighbor (WNN) analysis, an unsupervised method that evaluates the contribution of each modality and constructs a combined graph representing both RNA and ATAC data. WNN analysis was conducted using the FindMultiModalNeighbors function with 20 neighbors (k.nn), based on UMAP reductions obtained after Harmony^25^ integration. The WNN graph was then employed to generate a joint UMAP visualization. Clustering based on the WNN graph was performed using the FindClusters function with the smart local moving (SLM) algorithm. Clusters with fewer than 50 nuclei, a high percentage of doublets, or significant enrichment in mitochondrial markers were excluded from further analysis.

### Cluster annotation

Cell type annotation was performed using Azimuth^20^, with the Allen Human Motor Cortex Atlas^31^ as the reference dataset. Azimuth performed label transfer by assigning each cell in the MOGHE dataset to the most similar cell type in the reference based on gene expression similarity. The cell type assigned to each cluster was determined by identifying the cell type with the highest mean frequency within that cluster. The Azimuth annotation was manually validated using marker genes, as depicted in Figure 1E. Cluster markers were identified using the FindAllMarkers function on normalized data. Differentially expressed cluster markers were determined with the Wilcox rank-sum test, applying a log2 fold-change threshold of 0.25 and an adjusted P value of less than 0.05. For the oligodendrocyte subcluster analysis, markers were determined using the Wilcox rank-sum test in Seurat with the following parameters: log2 fold-change > 1, percent of cells expressing the marker (pct.1) > 0.3, and adjusted P < 0.05. Mitochondrial genes and those located in sex chromosomes were removed from marker genes.

**Figure 1.**
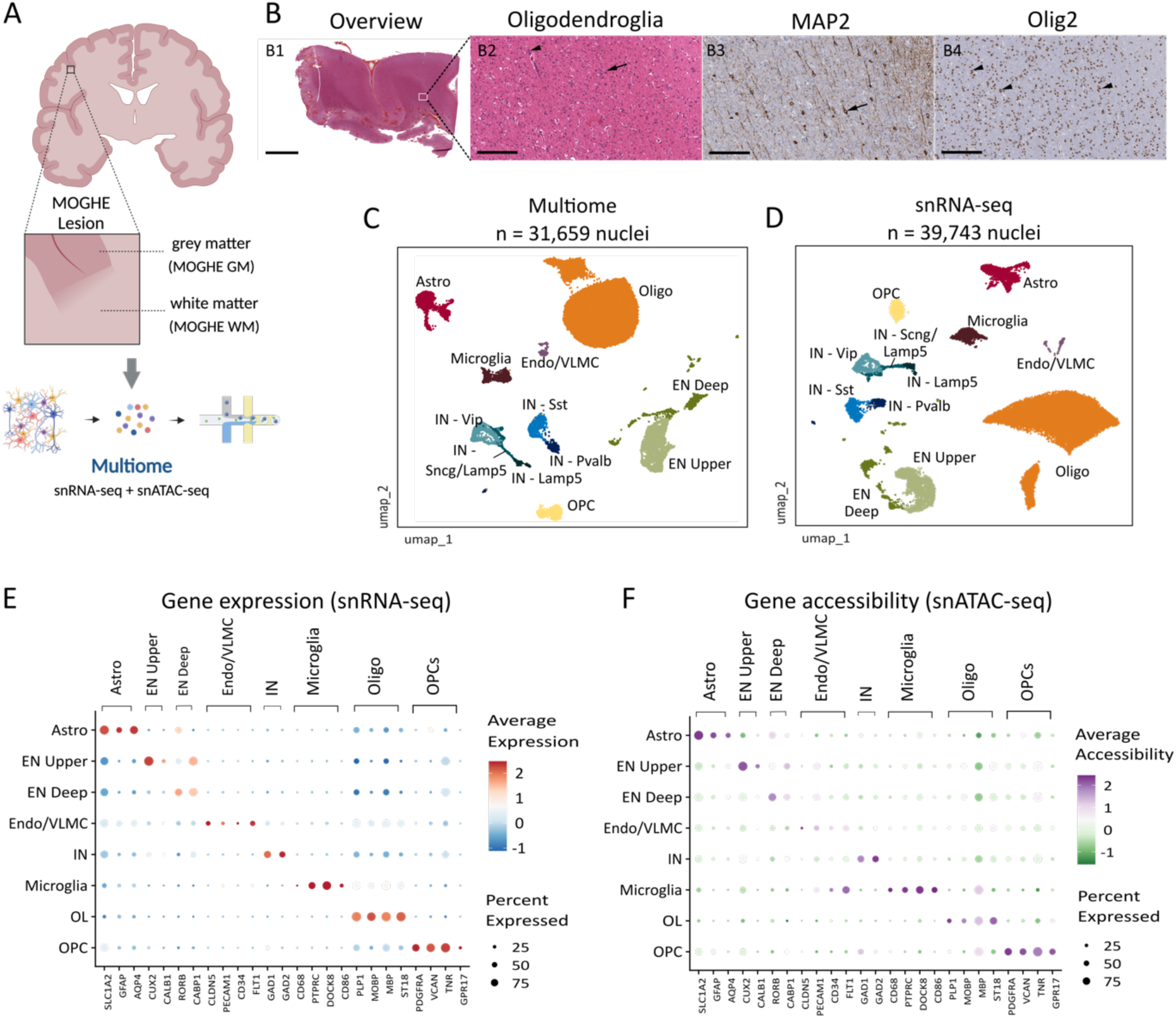
Multimodal single-nucleus sequencing of MOGHE tissue. **(A)** Schematic of MOGHE samples and study design. Nuclei from the MOGHE grey matter (MOGHE-GM) and white matter regions (MOGHE-WM) isolated and profiled by snRNA-seq and snATAC-seq using the Multiome assay (10X Genomics). Created with Biorender. **(B)** Representative histopathological findings of a brain surgical sample with MOGHE sequenced in the study. **B1.** Hematoxylin and eosin (H&E) stained section showing an overview of the resected tissue. The white rectangle indicates a subcortical WM region near the GM-WM junction presented in B2-B4. **B2.** Clusters of oligodendroglial cells either perivascular (arrow head) or around heterotopic neurons (arrow). **B3.** Immunohistochemistry (IHC) for MAP2 (neurons) depicts scattered heterotopic neurons (arrow). **B4.** IHC for Olig2 (oligodendrocytes) showing an irregularly distributed increase of the oligodendroglial cells, either isolated or in clusters (arrow heads). Scale bars: 4 mm (Figure B1) and 200 µm (Figures B2-B4). **(C)** Joint uniform manifold approximation and projection (UMAP) visualization of snATAC-seq and snRNA-seq from nuclei sequenced in MOGHE samples, colored by annotated cell type. **(D)** UMAP visualization of snRNA-seq from MOGHE samples integrated with neurotypical WM controls obtained from Elkjaer et al.^29^, and colored by cell type. **(E)** Dot plot displaying gene expression levels for canonical markers of major cortical brain cell types. The size of each dot represents the proportion of cells expressing the marker, while the color indicates the average expression levels. **(F)** Dot plot displaying gene accessibility levels computed on the basis of chromatin accessibility for the same marker genes shown in (E). The size of each dot represents the proportion of cells expressing the marker, while the color indicates the average gene accessibility. Astro, astrocytes. EN, excitatory neurons. Endo/VLMC, endothelial/vascular and leptomeningeal cells. IN, inhibitory neurons. OL, oligodendrocytes. OPC, oligodendrocyte precursor cells.

### Differential cellular abundance

We utilized the permutation_test function from the scProportionTest R package^32^ v. 0.0.0.9 to assess changes in cell type composition between tissue types (MOGHE-WM *vs* Control-WM and MOGHE-WM *vs* MOGHE-GM). The significance test used 10,000 permutations, and cell type changes with a false discovery rate (FDR) less than 0.05 and a fold difference greater than 2 were considered statistically significant.

### Gene signature analysis

Gene signature scores were computed using the AddModuleScore function from the Seurat package applied to log-normalized RNA data. Signatures for oligodendrocyte subtypes were obtained from Jakel et al., 2019^33^ and Sadick et al., 2022^34^.

### Motif enrichment analysis

We utilized chromVAR v3.3.2^35^ and Signac to perform motif analysis. Nuclei-level motif activity was computed using the RunChromVAR function in Signac with the set of background peaks matched appropriately. Differential activity of motifs was determined using the FindMarkers function, applying a likelihood ratio (LR) test with ATAC fragment counts as a latent variable, a minimum percentage of cells expressing the motif (min.pct) set to 0.05, and an adjusted p-value cutoff of < 0.05. To ensure biologically-relevant results, motifs were further filtered based on the expression of their associated regulators, which were required to be expressed in at least 10% of the cells within a given cluster/cell type.

### Cellular communication analysis

Cellular communication was performed using the CellChat R package v.2.1.2^36^. In brief, CellChat utilizes a mass-action-based model to quantify the signaling communication probability between two cell groups using a database of known ligand-receptor interactions. The cellular communication probabilities were computed using the computeCommunProb function with the parameter type set to triMean, and communications between groups with fewer than 10 cells were filtered out. Differential signaling interactions between MOGHE subclusters were calculated by performing differential expression analysis with identifyOverExpressedGenes and netMappingDEG functions, with a percentage of cells expressing the gene > 0.1 and p-value < 0.05. Significant interactions were classified as upregulated for ligands exhibiting a fold-change > 1.5.

## RESULTS

### Multimodal single-nucleus profiling of MOGHE lesions

We obtained frozen brain specimens from two donors aged 22 and 57 years with a confirmed pathological diagnosis of MOGHE after epilepsy surgery (Table 1). After isolating nuclei from the MOGHE gray matter (MOGHE-GM) and white matter regions (MOGHE-WM), we performed paired snRNA-seq and snATAC-seq on each nucleus using the 10X Genomics Multiome assay to measure chromatin accessibility and gene expression at nuclei resolution (Figure 1A). The sequenced tissues fulfilled the histopathological criteria for MOGHE, presenting with increase of the oligodendrocyte density in the WM and excess of heterotopic neurons (Figure 1B, Methods). Deep exome sequencing of the donors using the MOGHE-WM has not detected somatic variants in *SLC35A2* or in genes previously reported in epileptogenic brain lesions^37^.

**Table 1.**
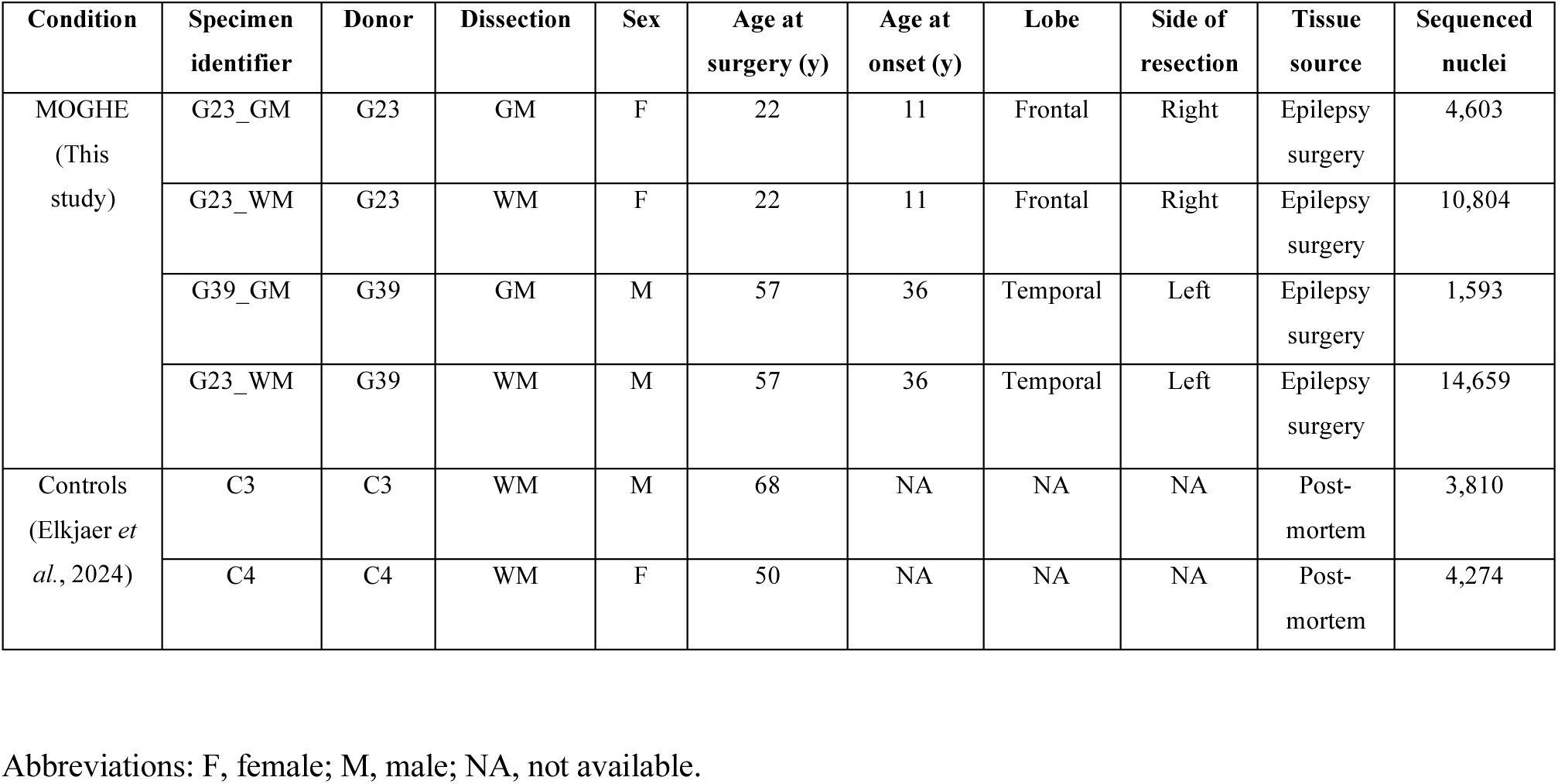
Description of the clinical samples.

After filtering to remove low-quality nuclei and doublets (Methods), we obtained 31,659 nuclei derived from MOGHE samples in the multiome snATAC-seq + snRNA-seq dataset (Figure 1C). Next, we integrated our MOGHE dataset with snRNA-seq performed on white matter from two neurotypical individuals (Control-WM) obtained from Elkjaer et al.^29^, resulting in a total of 39,743 nuclei (Figure 1D, Table 1). Clustering analysis identified 33 distinct clusters after integration of MOGHE and control nuclei (Figure S1A). Cell type annotation was carried out using Azimuth^20^, and subsequently validated through manual inspection of canonical marker genes (Figure 1E). The Azimuth annotation enabled confident labeling of most of these clusters (Figure S1B), and most clusters were shared across multiple samples (Figure S1C). After annotation, the majority of nuclei was classified as oligodendrocytes (OLs), followed by excitatory neurons (ENs, further divided into upper-and deep-layer subsets), astrocytes, inhibitory neurons (INs, subclassified into Vip, Sst, Pvalb, Lamp5 and Scng subtypes), oligodendrocyte precursor cells (OPCs), microglia and endothelial/vascular and leptomeningeal cells (Endo/VLMCs). As expected, samples derived from white matter (MOGHE-WM and Controls-WM) were predominantly composed of OLs, while neurons were not detected in Controls-WM and primarily located within MOGHE-GM (Figure S1D).

We also examined gene accessibility profiles computed based on the chromatin accessibility (snATAC-seq) for the same set of markers (Figure 1F). Gene accessibility displayed a strong correlation with gene expression across cell type markers, with the exception of some endothelial markers, denoting the agreement between data modalities. Thus, our single-nucleus profiling identified the various cortical cell types in GM and subcortical WM regions, revealing the cellular diversity of MOGHE lesions.

### Cellular and gene expression changes in MOGHE white matter

To investigate disease-associated changes in the WM, we analyzed cell type proportions between MOGHE-WM and Controls-WM as well as MOGHE-WM and MOGHE-GM. Compared to Controls-WM, MOGHE-WM tissue had a higher proportion of ENs, INs and microglia, denoting the expansion of heterotopic neurons and OLs in the WM of the abnormal tissue (Figure 2A). Also, MOGHE-WM had an increased proportion of OLs and a decreased abundance of astrocytes, ENs, and INs compared to MOGHE-GM, reflecting the expected differences in cell composition between cortical and subcortical regions (Figure 2B).

**Figure 2.**
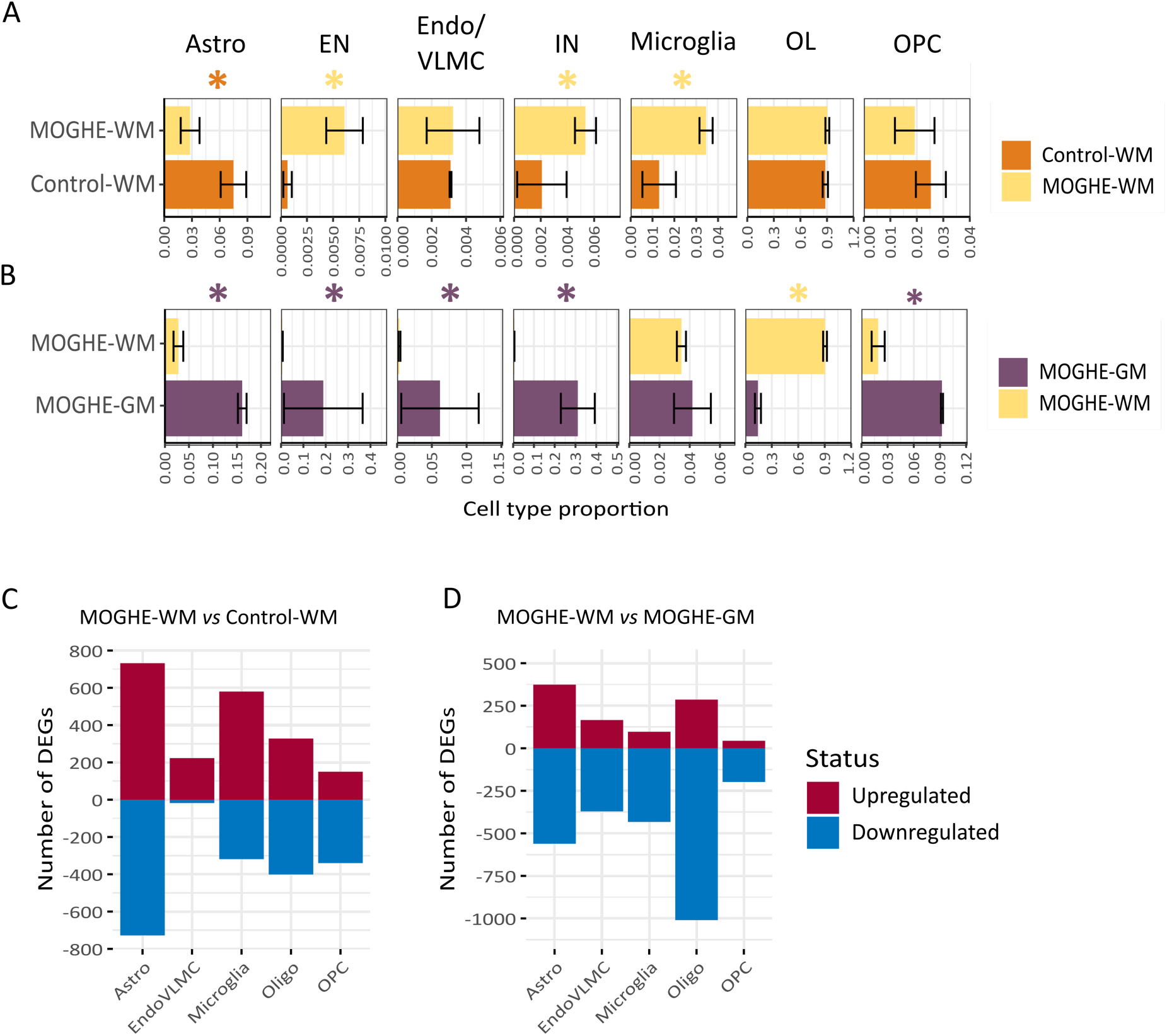
Cell type abundance and gene expression changes in MOGHE. **(A)** Barplots denoting cell types abundance in MOGHE-WM and Control-WM samples. The *x*-axis indicates the cell type proportion, and the *y*-axis indicates the tissue type. Error bars indicate the standard error of the mean. Differential cell type abundance was performed using the scProportionTest tool (Methods). Asterisks indicate significant changes considering an FDR < 0.05 and fold-change > 1.5. The asterisk color indicates the tissue type with the higher abundance. **(B)** Barplots denoting cell types abundance in MOGHE-WM and MOGHE-GM samples. Legends are defined as in (A). **(C)** Differentially expressed genes (DEGs) between MOGHE-WM and Control-WM, in non-neuronal populations. DEGs were obtained using MAST, with a fold-change > 2 and adjusted P value < 0.05. (see Methods). The *x*-axis indicates the cell type, and the *y*-axis indicates the number of DEGs. **(D)** DEGs in MOGHE-WM vs MOGHE-GM. Legend is defined in (C). Astro, astrocytes. EN, excitatory neurons. Endo/VLMC, endothelial/vascular and leptomeningeal cells. IN, inhibitory neurons. OL, oligodendrocytes. OPC, oligodendrocyte precursor cells.

We also analyzed differentially expressed genes (DEGs) between tissue conditions and within each cell type. Neurons were not considered in this analysis since they were not identified in Controls-WM. In MOGHE-WM compared to Controls-WM, there were significant gene expression changes in most glial populations, including astrocytes, microglia, and OLs (Figure 2C). When compared to MOGHE-GM, OLs in MOGHE-WM were the most affected population with a large number of downregulated genes (Figure 2C). Collectively, these analyses indicate that the cellular composition of subcortical MOGHE-WM was characterized by an increased OL population and the presence of heterotopic neurons, which recapitulated key hallmarks of the disease. Meanwhile, gene expression changes revealed that nearly all glial cell types in subcortical MOGHE-WM were affected by the condition.

### Pathological oligodendrocyte states in MOGHE

To identify MOGHE-associated OLs, we performed a subclustering analysis in the OLs compartment and assessed whether these subpopulations were enriched in disease tissue. Notably, subclusters 4 and 7 (OL_4 and OL_7), which clustered separately from the major OLs population, were predominantly observed in MOGHE (Figure 3A). Differential cellular abundance analysis confirmed that OL_4 and OL_7 were significantly expanded in MOGHE-WM compared to Controls-WM (Figure 3B, top). We inspected the subcluster distribution in individual samples and according to their localization (Figures S2A,B). The OL_4 subcluster was detected in both sequenced MOGHE samples and derived from WM and GM regions (n = 2,344 cells, 48% MOGHE-WM, 52% MOGHE-GM). The OL_7 subcluster was detected mostly in the WM of the G23 sample (n = 1,201 cells, 97% MOGHE-WM, 3% MOGHE-GM). Thus, this analysis uncovered disease-specific OLs subpopulations emerging in the cortical and subcortical regions of MOGHE lesions.

**Figure 3.**
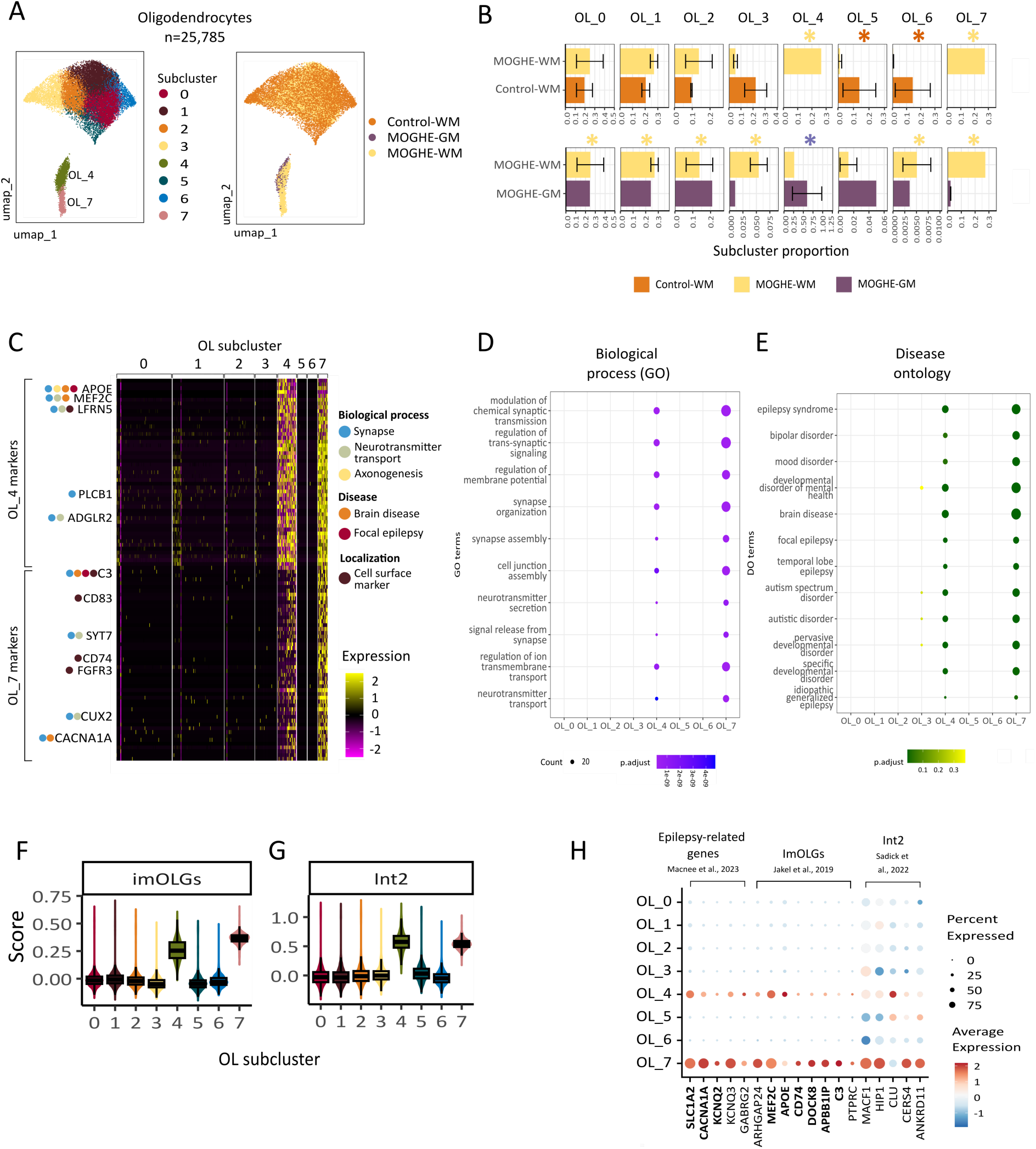
Identification of a MOGHE-specific oligodendrocyte population. **(A)** UMAP representation of oligodendrocytes (OLs) from MOGHE-WM, MOGHE-GM and Control-WM, colored by subcluster (left) and tissue type (right). **(B)** Bar plot showing the OL subcluster proportions in MOGHE-WM and Control-WM samples. The *x*-axis indicates the subcluster proportion, and the *y*-axis indicates the tissue type. Differential subcluster abundance was performed using the scProportionTest tool (Methods). Asterisks indicate significant changes considering an FDR < 0.05 and fold-change > 2. The asterisk color indicates the tissue type with the higher abundance. Error bars indicate standard error of the mean. **(C)** Heatmap showing top marker genes of MOGHE-specific OL_4 and OL_7 subclusters. Biological functions or disease association of selected markers are indicated. The complete list of markers is provided in Supplemental Table 1. **(D)** Dot plot showing top enriched Gene Ontology (GO) terms in OL subclusters computed using clusterProfiler^26^. The size of each dot represents the number of genes associated with the GO term, while the color reflects the enrichment P-value. **(E)** Dot plot showing top enriched Disease Ontology (DO) terms in OL subclusters computed using DOSE^27^ and clusterProfiler^26^. The size of each dot represents the number of genes associated with the DO term, while the color reflects the enrichment P-value. **(F)** Boxplots depicting activity scores of the immune oligodendroglia (ImOLG) signature from Jakel et al^33^ in OL subclusters. The *x*-axis indicates the subcluster, and the *y*-axis indicates the activity score. The center line of the boxplot shows the median of the data; the box limits show the upper and lower quartiles; the whiskers show 1.5 times interquartile ranges. Overlay dots represent activity scores in individual nuclei. **(G)** Boxplots depicting activity scores of the Int2 OL signature from Sadick et al^34^ in subclusters. Legends are defined in (H). **(H)** Dot plot displaying expression levels of gene sets overexpressed in OL_4 and OL_7: epilepsy-associated genes from Macnee et al.^38^, genes from the ImOLG signature from Jakel et al.^33^, genes from the Int2 signature from Sadick et al.^34^, and disease-associated oligodendrocytes (DAOs) from Pandey et al.^64^. The size of each dot represents the proportion of cells expressing the gene in subclusters, while the color indicates the normalized expression level.

Next, we used two approaches to functionally annotate these MOGHE-associated OLs. First, we performed differential marker analysis followed by enrichment analysis to identify subcluster-enriched biological themes. This approach revealed subcluster-specific marker genes for OL_4 (n = 220, adj. P < 0.05) and OL_7 subclusters (n = 967, adj. P < 0.05) (Supplemental Table 1, top 50 markers shown in Fig. 3C). OL_4 marker genes were strongly associated with synapse-related functions such as modulation of chemical synaptic transmission (adj. P = 6.6 × 10^-19^), regulation of trans-synaptic signaling (adj. P = 6.6 × 10^-19^), regulation of membrane potential (adj. P = 1.6 × 10^-15^), synaptic organization (adj. P = 4.9 × 10^-14^) and synapse assembly (adj. P = 4.7 × 10^-10^) (Figure 3D, Supplemental Table 1). Notably, these synaptic functions were also overrepresented among OL_7 markers, however they were not enriched in other OLs subclusters (Figure 3D). For instance, top expressed OL_4 markers included synaptic-related genes *APOE4*, *MEF2C*, *LFRN5*, *PLCB1*, *ADGLR2* (Figure 3C). On the other hand, top OL_7 markers included synaptic-related genes *C3*, *SYT7*, *CUX2* and *CACNA1A* (Figure 3C).

Based on these observations, we used the Synaptic Gene Ontology (SynGO), a comprehensive resource containing curated annotation of synaptic genes, to further investigate the functions associated with OL_4 and OL_7 subsets. In the OL_4 subcluster, we found that 28 genes (out of top 50 markers) were linked to synaptic components (q = 6.8 × 10^-18^, Figure S2C, Supplemental Table 2), of which 17 were postsynaptic genes (q = 2 × 10^-10^) and 15 presynaptic genes (q = 4.1 × 10^-10^). Similarly, we observed significant enrichment for synaptic genes in OL_7 (n = 13, q = 1.5 × 10^-4^), especially posynaptic genes (n=10, q = 1.7 × 10^-4^) (Figure S2C, Supplemental Table 2). Of note, functional enrichment using the Disease Ontology showed that MOGHE-specific OL_4 and OL7 were the only subclusters enriched for genes related to epilepsy and brain diseases (Figure 3E, Supplemental Table 1). For instance, OL_4 and OL_7 expressed markers linked to epilepsy^38^ such as *SLC1A2*, *KNCQ2*, *KCNQ3* (Figure 3H). Collectively, these analyses showed that MOGHE-specific OLs upregulated synaptic components and epilepsy-related genes.

The other approach to characterize MOGHE OLs involved comparison with gene signatures from pathological OLs in the literature. Previous snRNA-seq studies described the OLs subset known as the immune subtype (ImOLGs) and the integrated 2 subtype (Int2), which were identified in multiple sclerosis and Alzheimer’s disease, respectively^33,34^. We used the AddModuleScore from Seurat to score these pathological signatures in OLs identified in MOGHE lesions. Notably, OL_4 and OL_7 exhibited the highest scores for ImOLGs (Figure 3F) and Int2 (Figure 3G) signatures, indicating that MOGHE-specific OL_4 and OL_7 shared marker genes with these OLs subtypes identified in other neurological diseases. Common markers with the ImOLGs subtype included *MEF2C* and *APOE* (expressed in OL_4), and immune genes *CD74*, *DOCK8*, *APBB1IP* and *C3*, which are highly expressed in OL_7 (Figure 3H).

In addition, *OLIG2*, which encodes the protein considered to be a neuropathological marker of the expanded oligodendroglia cell clusters in MOGHE^2,7,39^, was expressed only in the MOGHE-specific OL_7 subpopulation (Figure S2E), even though the chromatin in *OLIG2* promoter region was accessible across all OL subclusters (Figure S2F). On the other hand, *OLIG2* was not detected in MOGHE-specific OL_4, thus indicating that other markers need to be considered for detecting pathological OLs in the tissue.

Altogether, these in-depth analyses of the OL compartment enabled us to identify MOGHE-specific OLs cell states associated with synapse regulation and sharing common markers with pathological OLs from other neurological conditions.

### Cellular communication of MOGHE oligondendrocytes

We inferred the cell communication network among the identified OLs subclusters and other cell types using CellChat^40^. First, we analyzed the global communication patterns by considering major cell types and including OL subclusters (Figure 4A). Neurons exhibited the most robust communication patterns, particularly with other neurons. Notably, among the OL subclusters, OL_4 and OL_7 showed the highest communication strength, especially with neurons and OPCs.

**Figure 4.**
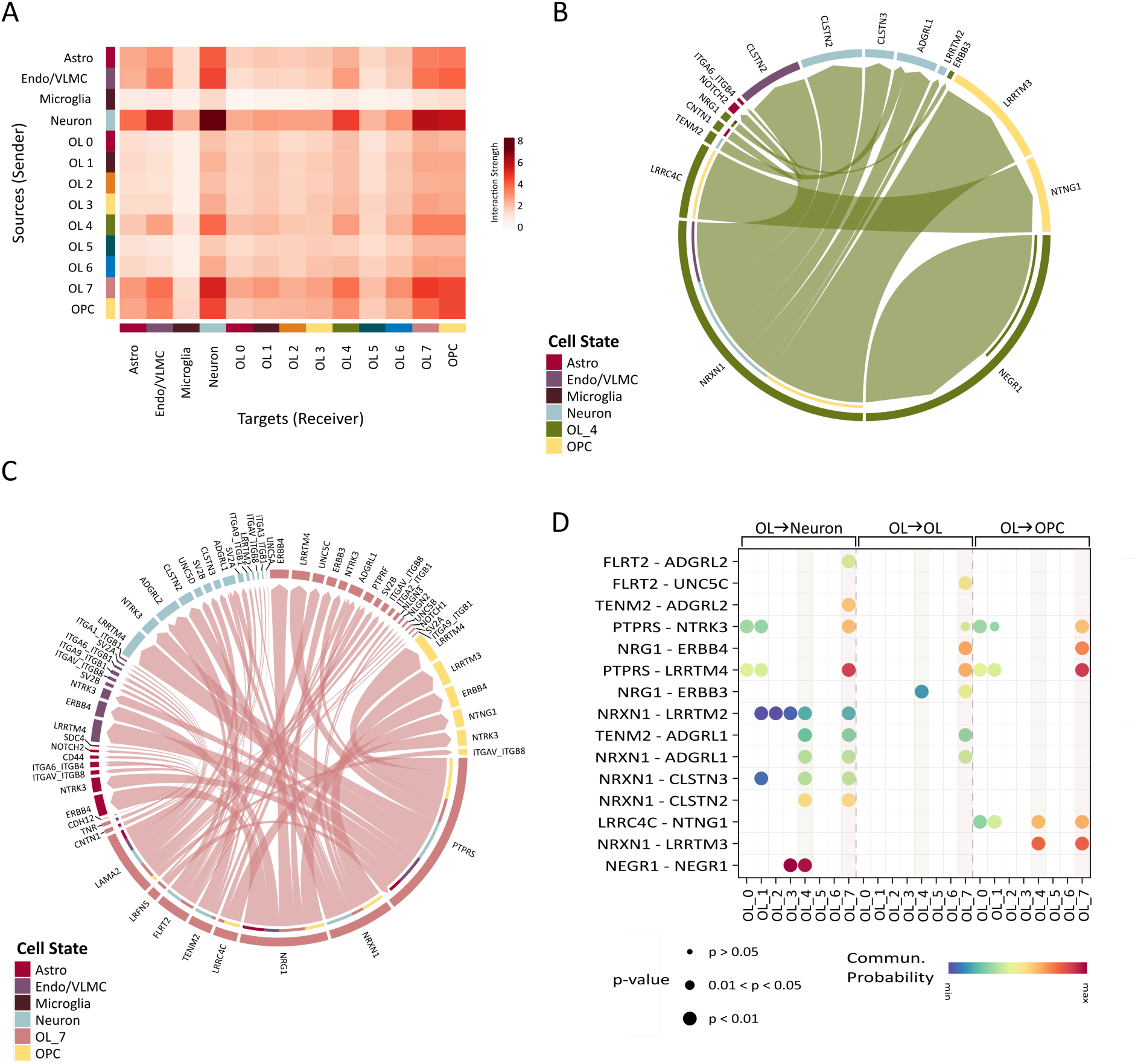
Cell communication analysis of oligodendrocytes (OLs) subclusters. **(A)** Heatmap showing interactions across cell types and OLs subclusters inferred by CellChat. The *y*-axis indicates the sources (senders) of the interaction, and the *x*-axis indicates the targets (receivers) of the interaction. Color denotes the interaction strength. **(B)** Chord diagram of upregulated ligand-receptor pairs in OL_4 compared to remaining OLs. Colors indicate cell states. Chords are drawn from ligands to receptors and the chord width is proportional to interaction strength. Inner color bars indicate the targets of the corresponding outer color bar. Significant interactions were computed based on percent of cells expressing the ligand > 0.1, p-value < 0.05, and ligand fold-change > 1.5. **(C)** Chord diagram of upregulated ligand-receptor pairs in OL_7 vs remaining OLs. Legends are defined in (B). **(D)** Dot plot of 15 top ligand-receptor pairs upregulated in OL 4 and OL 7 with highest communication probability. The *x*-axis indicates the OL subcluster, and the *y*-axis indicates the ligand-receptor pairs. Dot size indicates the p-value, and dot color indicates the interaction strength. Interactions are shown with OL as sources and either neurons (left), OLs (middle), or OPCs (right) as targets. Light gray vertical bars highlight OL_4 and OL_7 interactions.

Next, we conducted a differential analysis using CellChat to identify specific signaling interactions in OL_4 and OL_7 compared to other OL subclusters. This analysis revealed 12 and 63 significantly upregulated ligand-receptor pairs in OL_4 and OL_7, respectively. Examples of upregulated interactions in OL_4 (Figures 4B,D) included neuronal growth factor signaling (NEGR1 (OL) → NEGR1 (OL/Neuron)), neurexin signaling (NRXN1 (OL) → CLSTN2 (Neuron) and NRXN1 (OL) → LRRTM3 (OPC)), and neuroligin signaling (LRRC4C (OL) → NTNG1 (OPC)). In OL 7 (Figures 4C,D), the upregulated pathways involved PTPR signaling (PTPRS (OL) → LRNF5 (OL) and PTPRS (OL) → LRRTM4 (Neuron/OPC)), neurexin signaling (NRXN1 (OL) → NGLN3 (OL) and NRXN1 (OL) → CLSTN2 (Neuron)), and neuregulin signaling (NRG1 (OL) → ERBB3 (OPC/OL)). In line with previous GO analyses, these interactions within the neurexin, neuroligin and neuregulin signaling pathways are implicated in synaptic organization^41,42^ and oligodendrocyte differentiation^43,44^. These results highlight the distinct communication patterns of MOGHE-specific OL_4 and OL_7 subclusters particularly through interactions with neurons and OPCs, suggesting their functional specialization in synaptic regulation.

### Characterization of heterotopic neurons in MOGHE

Next, given that the appearance of heterotopic neurons (HNs) is a key histological feature of MOGHE, we asked whether misplaced neurons could be identified in the WM of MOGHE samples. We identified a subpopulation of HNs derived specifically from MOGHE-WM in both patients (n = 246; Figures 5A,B), while HNs were not detected in Control-WM samples. Most of these HNs were annotated as ENs from layers 5/6 (n = 129; 52%), and the remaining nuclei consisted mainly of INs subtypes Lamp5, Sst and PvalB. The HNs subpopulation expressed a very distinct set of markers such as cell-adhesion genes (*CDH5* and *CDH3*), transmembrane proteins (*TMEM275* and *TMEM233*), ion channels (*KCNK2*, *CACNA1B*), and the neurotransmitter *NPY* (Figure 5C, Supplemental Table 3).

**Figure 5.**
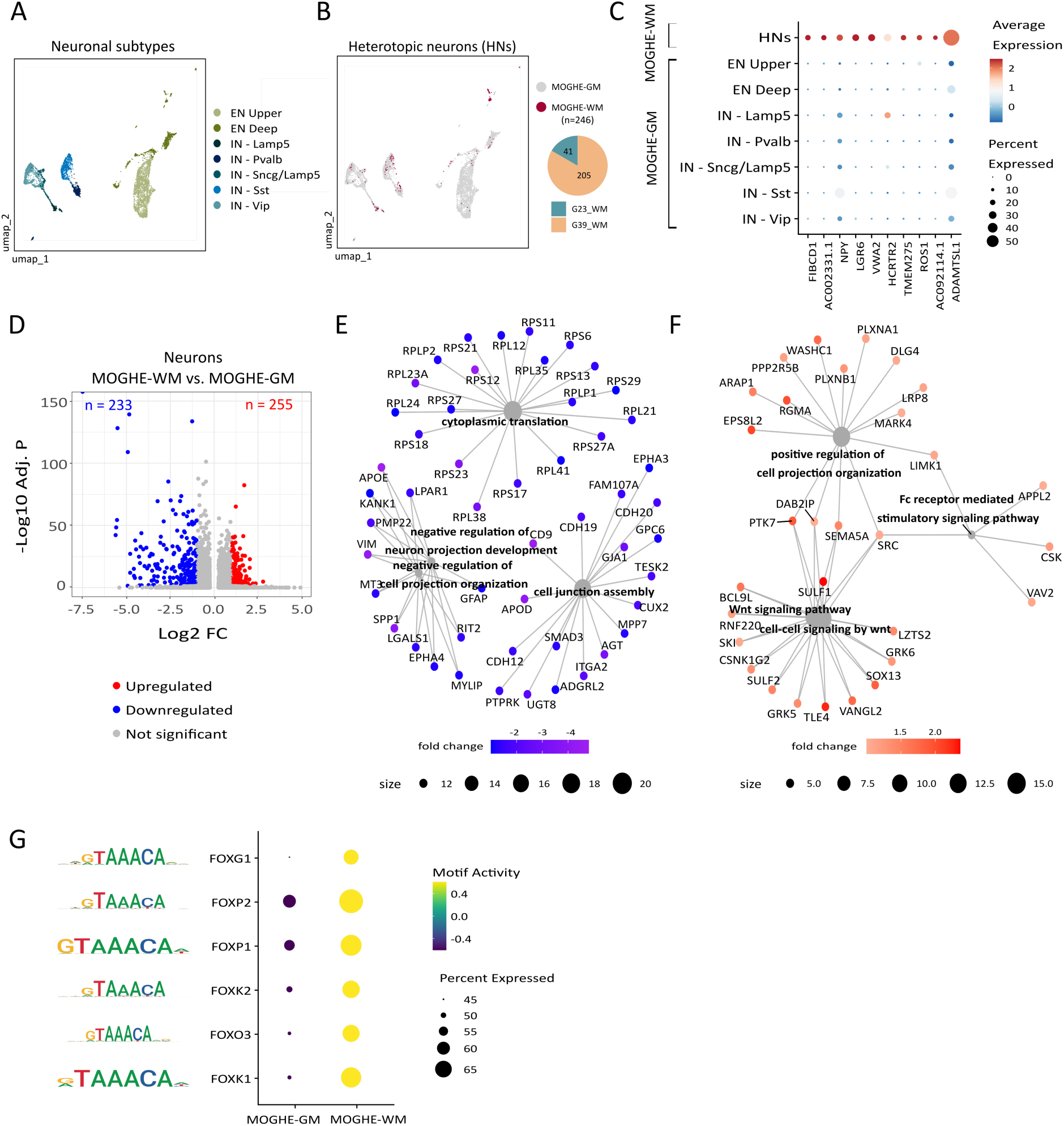
Characterization of heterotopic neurons in MOGHE. **(A)** UMAP representation of neuronal nuclei from MOGHE-WM and MOGHE-GM, colored by neuronal subtype. **(B)** UMAP representation of neuronal nuclei from (A), colored by tissue origin. Heterotopic neurons (HNs) are indicated in red. **(C)** Dot plot displaying gene expression of HNs marker genes. The size of each dot represents the proportion of cells expressing the marker, while the color indicates the average expression levels. **(D)** Volcano plot displaying differentially expressed genes (DEGs) in MOGHE-WM *vs* MOGHE-GM. DEGs were obtained using MAST, with a fold-change > 2 and adjusted P value < 0.05. (see Methods). The *x*-axis indicates the log2 fold-, and the *y*-axis depicts the significance of the change (-log10 of the adjusted P-value). The color indicates the gene status. The complete list of DEGs is available on Supplemental Table 3. **(E)** Gene-term network depicting downregulated genes and corresponding Gene Ontology (GO) molecular functions. GO terms are represented as grey nodes, with size corresponding to the number of associated genes. Genes are colored according to the fold-change values. Enriched GO terms were computed with clusterProfiler. **(F)** Gene-term network illustrating upregulated genes and corresponding molecular functions. Legends are defined in (E). **(G)** Dot plots depicting motif activity of FOX transcription factors in MOGHE-GM and MOGHE-WM neurons. Motif activity was computed using chromVar based on the snATAC-seq datasets. Dot color corresponds to motif activity, and the size corresponds to the proportion of nuclei with open chromatin peaks containing a binding site for the motif. The motif sequence recognized by the regulator is indicated in the left.

To identify molecular functions in this abnormal neuronal subpopulation, we analyzed DEGs between HNs (i.e. MOGHE-WM neurons) and total MOGHE-GM neurons, identifying 255 upregulated and 233 downregulated genes (Figure 5D, Supplemental Table 3). Enriched GO terms associated with downregulated genes included negative regulation of neuron projection, cytoplasmic translation, and cell junction assembly (Figure 5E). On the other hand, upregulated molecular functions included positive regulation of cell projection, the Fc receptor-mediated stimulatory signaling pathway, and the Wnt signaling pathway (Figure 5F), which has been shown to regulate neuronal migration^45,46^.

In line with the DEG analysis, a motif enrichment analysis based on the snATAC-seq data identified that FOX transcriptional regulators were among the top activated motifs in HNs (Figure 5F). The FOX family has been shown to regulate the Wnt signaling pathway^47,48^, which could explain the distinctive Wnt activation in HNs. Therefore, these results indicate that HNs in MOGHE consisted of ENs from deep cortical layers with increased expression of pathways associated with neuronal migration.

## DISCUSSION

In this study, we applied multiomic single-nucleus profiling to systematically map the cellular landscapes of cortical and subcortical regions affected in MOGHE, a type of focal MCD characterized by white matter abnormalities and associated with epilepsy. We identified significant alterations in cell type composition in the WM of MOGHE tissues and uncovered distinct pathological cell populations in this condition.

We identified disease-specific OLs subpopulations emerging in MOGHE tissue with unique transcriptomic profiles. Gene ontology and cell-to-cell communication analyses revealed functional differences between MOGHE-specific OLs subclusters and the remaining oligodendrocytes, notably the upregulation of synaptic genes and enhanced neuron communication. Thus, we postulate that MOGHE-specific OLs may be involved in synaptic support and the mediation of glial-neuron interactions in the disease.

OPCs are the specific glial cells that directly form synapses with neurons^49^. Yet, oligodendroglia are a transcriptionally heterogeneous lineage with distinct populations emerging in response to neurological conditions^50,33,51^. For example, snRNA-seq in human OLs from WM controls and multiple sclerosis found that OPCs and committed oligodendrocytes precursors (COPs) were enriched in synapse regulation^33^. Sadick et al also identified OLs populations enriched for transcripts involved in synapse assembly and organization in Alzheimer’s disease^51^. Thus, the emergence of synapse-enriched OLs in neurological conditions associated with WM degradation and myelin loss^49,52^ such as multiple sclerosis and Alzheimer’s has been described, however the roles of these synapse-enriched OLs remain unknown.

Some of the markers associated with MOGHE OLs have established functions. For example, *APOE4*, a marker of the OL_4 subpopulation, has been shown to impair oligodendrocyte differentiation^53^ and downregulate myelination via cholesterol dysregulation^54^. Cholesterol biosynthesis is essential for myelin biogenesis^55,56^. Thus, these dysregulated OLs states might contribute to WM matter degradation and the formation of irregular zones of hypomyelination typically observed in MOGHE-WM. On the other hand, the OL_7 subcluster expressed markers of immune cells such as *C3*, *CD74*, and *CD83*, therefore sharing a common transcription profile with the immune oligodendroglia (ImOLG) population described in multiple sclerosis^33^.

Other glial cell types may also contribute to the disease, as evidenced by microglia increase in MOGHE-WM. Microglia have been linked to epilepsy and their effects can be pro-or anti-inflammatory depending on the disease stage^57,58^. Moreover, resident microglia are required for maintaining myelin health and their dysregulation may be related to the emergence of pathological OL states^59^.

We also identified the appearance of misplaced neurons in the WM (HNs), another key pathological hallmark of MOGHE. Even though MOGHE HNs were a rare population, we could identify specific biological functions in this neuronal subset. We found that MOGHE HNs displayed increased expression of neuronal projection genes and Wnt signaling pathway. Interestingly, Wnt signaling has been linked to axonal/dendrite remodeling and neuronal plasticity^60,61^, and regulates neuronal migration by modulating the microtubule cytoskeleton^45,46^, providing a potential mechanism for the abnormal localization of HNs. Furthermore, alterations in Wnt signaling induce neuronal hyperexcitability^62,63^, a defining feature of epilepsy. Thus, MOGHE HNs represent a disease-specific neuronal population that may contribute to the development of epileptogenic circuits in MOGHE.

While our findings provide valuable insights into the cellular landscape of MOGHE, this study has limitations due to its restricted sample size. For instance, while the MOGHE-specific OL_4 subcluster was observed in both patients, OL_7 was predominantly derived from a single patient.

Future studies with larger cohorts are warranted to confirm the MOGHE-associated OL subpopulations identified here and may reveal additional cellular states linked to the disease. Furthermore, our results indicated that some MOGHE-specific OLs derived from the OL_4 subcluster were located in the GM, possibly due to the blurring of the gray-white matter junction in lesions. Spatial transcriptomics and/or in situ immunohistochemistry studies using the markers identified in this study will be needed to determine the localization of MOGHE-specific OLs in lesions. In addition, incorporating cases with *SLC35A2* loss-of-function variants will be important for understanding the impact of this common genetic variation on the cellular heterogeneity of the disease.

In summary, this high-resolution mapping of MOGHE lesions in clinical samples represents the first cellular atlas of human tissue affected by the disease, providing a comprehensive view of the perturbed cell populations and gene expression alterations involved in this pathology.

## Data Availability

Raw and processed sequencing datasets from paired snRNA-seq and snATAC-seq generated in this study were deposited at the Gene Expression Omnibus under accession code GSE284073.

## Author contributions

D.F.T.V. and F.R. designed the study. M.L. performed single-nucleus assays. I.C.G. and D.F.T.V. performed data analyses. C.L.Y., E.G., H.T., M.K.M.A., F.C., and I.L-C contributed to clinical sample collection. L.K., I.B. and F.R. performed histopathological analyses. I.C.G. and D.F.T.V. wrote the manuscript. All authors reviewed and approved the final version of the manuscript.

## Ethics approval and patient consent

We confirm that we have read the Journal’s position on issues involved in ethical publication and affirm that this report is consistent with those guidelines. This study was approved by the University of Campinas’ Research Ethics Committee (CAAE: 12112913.3.0000.5404), and written informed consent was obtained from patients or their legal guardians.

## SUPPLEMENTARY FIGURES

**Figure S1.**
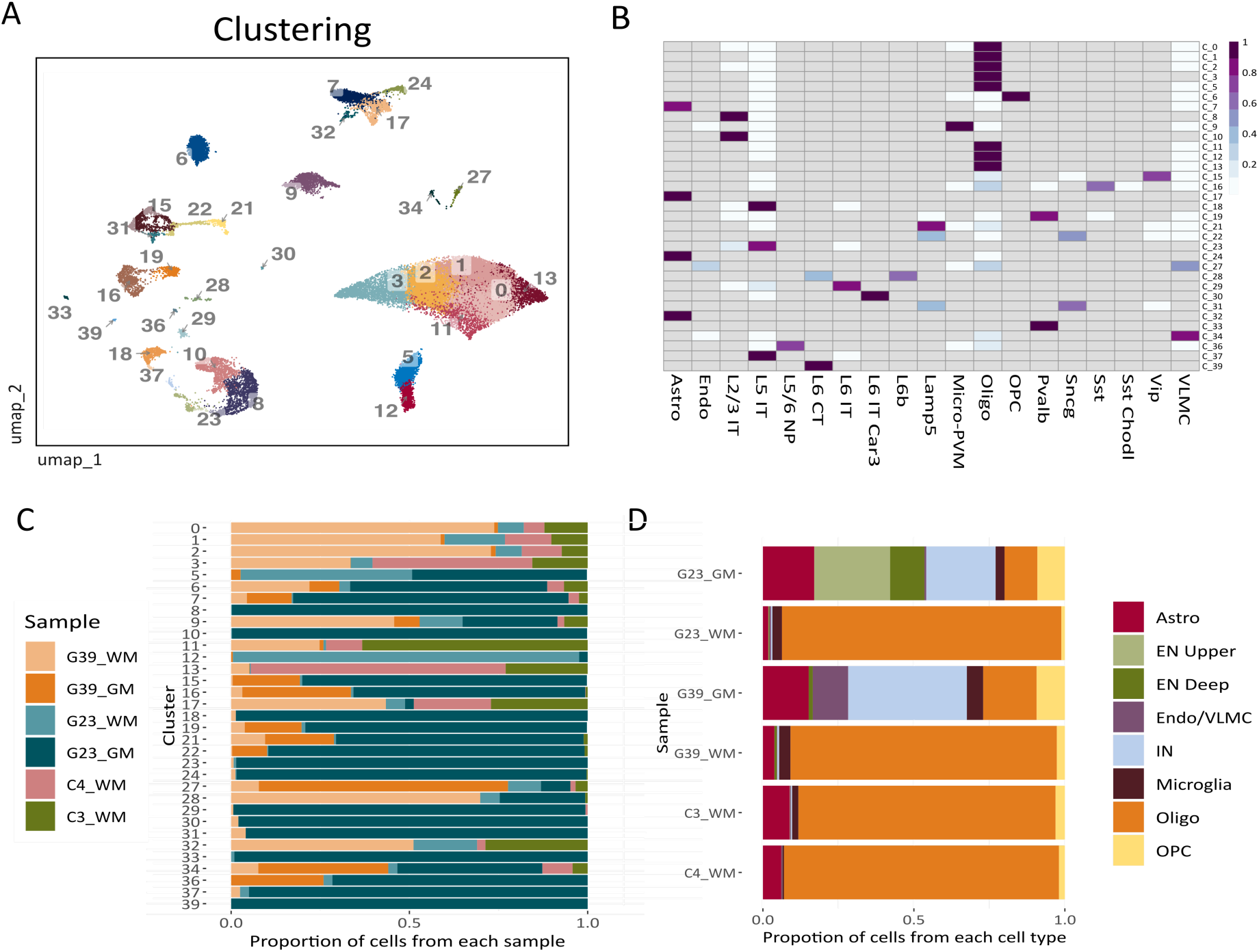
related to Figure 1. Clustering and annotation. **(A)** UMAP visualization of nuclei clusters. Clustering was performed in the RNA graph obtained after Harmony integration of MOGHE-WM, MOGHE-GM and Control-WM snRNA-seq samples. **(B)** Heatmap depicting the annotation of 40 nuclei clusters performed by Azimuth based on the Allen atlas of the human cortex as a reference. The color indicates the proportion of Azimuth subtypes identified in each cluster. **(C)** Bar graphs denoting the representation of samples across nuclei clusters. **(D)** Bar graphs showing the proportion of cell types across samples.

**Figure S2.**
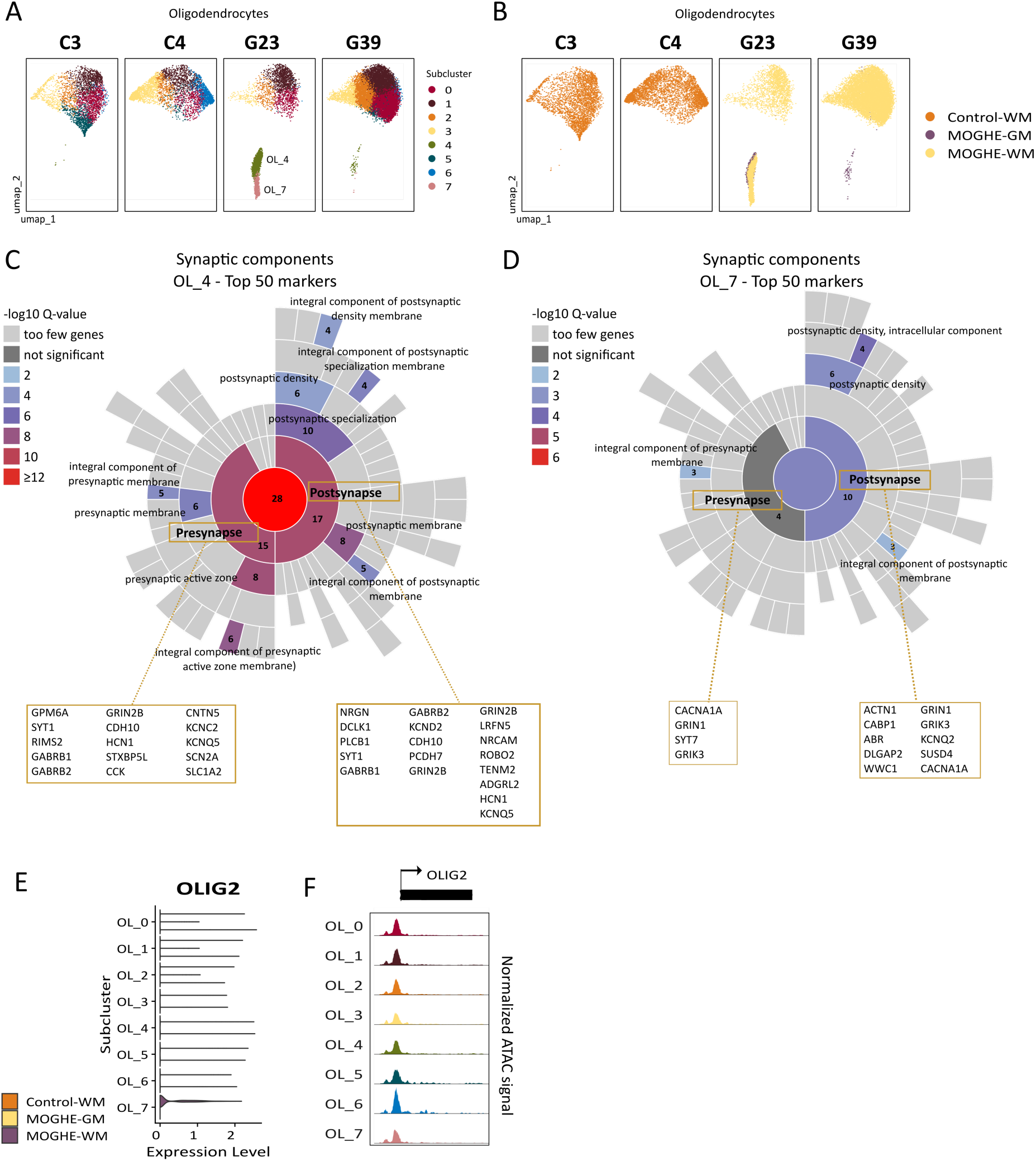
related to Figure 3 Additional information related to MOGHE-specific oligodendrocytes. (A) UMAP visualization of oligodendrocytes (OLs) in individual MOGHE samples (G23 and G29) and controls (C3, C4), colored by subcluster. **(B)** UMAP visualization of OLs in individual MOGHE samples and controls colored by tissue type. **(C)** Synaptic Gene Ontology (SynGO) term enrichment analysis based on the top 50 markers genes of the OL_4 subcluster. In the sunburst plot, terms are colored according to their Q-value, and the number of genes corresponding to each synaptic GO term is indicated. **(D)** SynGO term enrichment analysis based on the top 50 markers genes of the OL_7 subcluster. Legend is defined as in (C). **(D)** Violin plot of *OLIG2* expression in OL subclusters. The *x*-axis indicates the subcluster and the *y*-axis indicates the normalized expression level. **(E)** Chromatin accessibility profile of *OLIG2* in OL subclusters from snATAC-seq. The normalized ATAC signal is depicted in a region of +/− 1 kb around the gene start/end coordinates.

## SUPPLEMENTARY TABLES

**Supplemental Table 1.** Subcluster markers and Gene Ontology (GO)/Disease Ontology (DO) term enrichment analysis associated with OL_4 and OL_7 oligodendrocyte subclusters, related to Fig. 3.

**Supplemental Table 2.** Synaptic Gene Ontology (SynGO) term enrichment analysis based on the top 50 markers genes of OL_4 and OL_7 subclusters, related to Fig. S2.

**Supplemental Table 3.** Information related to heterotopic neurons including gene markers, differentially expressed genes, and enriched gene ontology terms, related to Fig. 5.

